# Understanding RNP remodelling uncovers RBPs functionally required for viral replication

**DOI:** 10.1101/350686

**Authors:** Manuel Garcia-Moreno, Marko Noerenberg, Shuai Ni, Aino I Järvelin, Esther González-Almela, Caroline Lenz, Marcel Bach-Pages, Victoria Cox, Rosario Avolio, Thomas Davis, Svenja Hester, Thibault J.M Sohier, Bingnan Li, Miguel A Sanz, Luis Carrasco, Emiliano P. Ricci, Vicent Pelechano, Bernd Fischer, Shabaz Mohammed, Alfredo Castello

**Affiliations:** Department of Biochemistry, University of Oxford, OX1 3QU Oxford, UK; Department of Chemistry, University of Oxford, Chemistry Research Laboratory, Mansfield Road, Oxford OX1 3TA, UK; German Cancer Research Center (DKFZ), 69120 Heidelberg, Germany; Faculty of Biosciences, Heidelberg University, Heidelberg, Germany; Centro de Biologia Molecular “Severo Ochoa”, Universidad Autonoma de Madrid, 28049 Madrid, Spain; Department of Molecular Medicine and Medical Biotechnology, University of Naples Federico II, Italy.; Université de Lyon, ENSL, UCBL, CNRS, INSERM, LBMC, 46 Allée d’Italie, 69007, Lyon, France.; SciLifeLab, Department of Microbiology, Tumor, and Cell Biology, Karolinska Institutet, 17165 Solna, Sweden

## Abstract

The compendium of RNA-binding proteins (RBPome) has been greatly expanded by the development of RNA-interactome capture (RNA-IC). However, it remains unknown how responsive is the RBPome and whether these responses are biologically relevant. To answer these questions, we created ‘comparative RNA-IC’ to analyse cells challenged with an RNA virus, called sindbis (SINV). Strikingly, the virus altered the activity of 245 RBPs, many of which were newly discovered by RNA-IC. Mechanistically, alterations in RNA binding upon SINV infection are caused by changes in the subcellular localisation of RBPs and RNA availability. Moreover, ‘RBPome’ responses are crucial, as perturbation of dynamic RBPs modulates the capacity of the virus to infect the cell. For example, ablation of XRN1 causes cells to be refractory to infection, while GEMIN5 moonlights as a novel antiviral factor. Therefore, RBPome remodelling provides a mechanism by which cells can extensively rewire gene expression in response to physiological cues.

**HIGHLIGHTS:**

- A quarter of the RBPome remodels upon SINV infection.
- The remodelling is caused by changes in protein localisation and RNA availability.
- Rewiring of the RBPome is crucial for viral infection efficacy.
- We discover RBPs with previously unknown anti- or pro-viral activity.

## INTRODUCTION

RNA-binding proteins (RBPs) assemble with RNA forming ribonucleoproteins (RNPs) that dictate RNA fate (Glisovic et al., 2008). Historically, most of the known RBPs were characterised by the presence of well-established RNA-binding domains (RBDs), which include the RNA recognition motif (RRM), K-homology domain (KH) and others (Lunde et al., 2007). However, stepwise identification of unconventional RBPs (e.g. (Chang et al., 2013; Hentze and Argos, 1991; Sampath et al., 2004)) evoked the existence of a broader universe of protein-RNA interactions than previously anticipated. Notably, *in silico* and *in cellulo* system-wide approaches have greatly expanded the compendium of RBPs (RBPome), adding hundreds of proteins lacking known RBDs and previous links with RNA biology (Baltz et al., 2012; Brannan et al., 2016; Castello et al., 2012; Gerstberger et al., 2014; Hentze et al., 2018). For example, RNA-interactome capture (RNA-IC) employs ultraviolet (UV) crosslinking, oligo(dT) capture under denaturing conditions and quantitative mass spectrometry to identify the repertoire of proteins interacting with polyadenylated [poly(A)] RNA in living cells (Baltz et al., 2012; Castello et al., 2012; Castello et al., 2013). Applied to HeLa cells, RNA-IC identified 860 RBPs, 315 of which were previously unrelated to RNA biology (Castello et al., 2012). Importantly, the crucial participation of several of these ‘unorthodox’ RBPs in RNA life has been proven in follow up studies (Hentze et al., 2018). How unorthodox RBPs interact with RNA remained, nevertheless, poorly understood. Several proteomic approaches tackled this question on a global scale, revealing that protein-protein interaction domains, enzymatic cores, DNA-binding domains and intrinsically disordered regions can function as RNA-binding surfaces (Castello et al., 2016; He et al., 2016; Kramer et al., 2014). However, it remains unknown whether the RBPome responds globally to physiological cues and if so, how these responses are triggered and what are their biological consequences. Recent work approached these questions by analysing the RBPome during fruit fly embryo development, and reported changes in RBP composition that could be explained by matching alterations in protein abundance. In other words, the proteome of the embryo changes through development and as a consequence the scope of available RBPs (Hentze et al., 2018; Sysoev et al., 2016). However, several RBPs did not follow this trend, displaying protein-level independent changes in RNA binding and raising the question of whether physiological perturbations can induce such responsive behaviour more widely. To address this possibility, we used, here, an improved ‘comparative RNA-IC’ approach to profile with high accuracy RBP dynamics in cells infected with sindbis virus (SINV) (Figure 1A and 1B).

**Figure 1.**
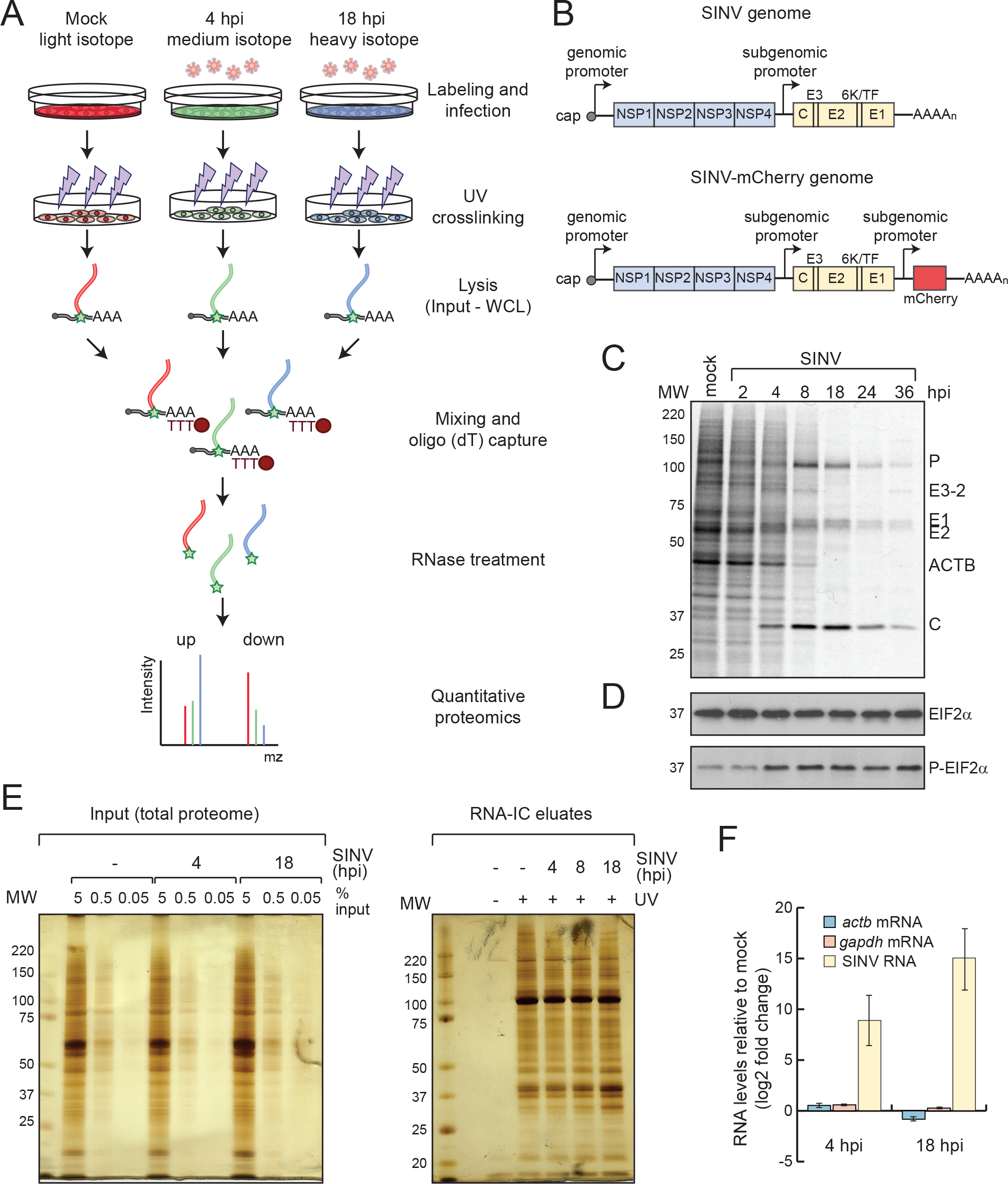
Implementation of RNA-IC to HEK293 cells infected with SINV. A) Schematic representation of RNA-IC combined with SILAC and virus infection. Several modifications were implemented into the original protocol (Castello et al., 2012; Castello et al., 2013), including the use of isotopic labelled amino acids and the combination of the samples after lysis and prior to the oligo(dT) capture. B) Schematic representation of SINV (wild type) and chimeric SINV-mCherry genomes. Subgenomic (sg)RNA is synthesised from the ‘subgenomic promoter’. C) Analysis of the proteins synthesised in uninfected and SINV-infected HEK293 cells by [^35^S]-Met/Cys labelling. D) Analysis of total and phosphorylated eIF2α in cells from panel (C) by western blotting. E) Silver staining analysis of the ‘inputs’ (i.e. total proteome, left) and eluates (i.e. RNA-bound proteome, right) of a representative RNA-IC experiment in SINV-infected HEK293 cells. F) RT-qPCR analysis of the eluates of RNA-IC experiments using specific primers against SINV RNAs, *gapdh* and *actb* mRNAs. Error bars represent standard deviation of three biological replicates. hpi, hours post infection. MW, molecular weight.

Viruses have been fundamental for the discovery and characterisation of important steps of cellular RNA metabolism such as RNA splicing, nuclear export and translation initiation. This is due to their ability to hijack the required host resources, often by interfering with the activity of the master regulators of these pathways (Berget et al., 1977; Castello et al., 2011; Castello et al., 2012; Chow et al., 1977; Krausslich et al., 1987; Lloyd, 2015; Sun and Baltimore, 1989; von Kobbe et al., 2000). Furthermore, specialised RBPs are at the frontline of cellular antiviral defences, detecting pathogen-associated molecular patterns (PAMPs) such as double stranded RNA or RNAs with 5’ tri-phosphate ends (Barbalat et al., 2011; Garcia et al., 2006; Rehwinkel and Reis e Sousa, 2013; Vladimer et al., 2014). Since RBPs are crucial for both the viral biological cycle and the host antiviral defences (Garcia-Moreno et al., 2018), virus infection thus represents an optimal scenario to assess the RBPome dynamics.

Our data show that the complement of active cellular RBPs is strongly altered in response to virus infection, revealing that the RBPome is highly dynamic. This phenomenon is not due to changes in protein abundance, but to an extensive subcellular redistribution of host RBPs and to alterations in the availability of RNA substrates. Importantly, RBP responses are biologically crucial, as dynamic RBPs modulate the viral replication cycle or/and the ability of the cell to counteract the infection. We envision that these proteins may represent novel targets for host-based antiviral therapies.

## RESULTS AND DISCUSSION

### Applying RNA-IC to cells infected with SINV

To study the dynamics of cellular RBPs in response to physiological cues, we challenged cells with a cytoplasmic RNA virus and applied RNA-IC. We chose SINV and HEK293 cells as the viral and cellular model, respectively. SINV is a highly tractable viral model that is transmitted from mosquito to vertebrates, causing high fever, arthralgia, malaise and rash in humans. Its biological cycle is relatively well-understood, involving a similar scope of cellular factors to other pathogenic alphaviruses such as chikungunya virus (CHIKV) and venezuelan equine encephalitis virus (VEEV) (Carrasco et al., 2018). SINV replicates in the cytoplasm of the infected cell and produces three viral RNAs (Figure 1B and S1A): the genomic (g) RNA; the subgenomic (sg)RNA and the negative stranded RNA. gRNA is packaged into the viral capsid and is translated to produce the non-structural proteins (NSP) that form the replication complex. The sgRNA is synthesised from an internal promoter and encodes the structural proteins (SP) required to generate the viral capsids. The negative strand serves as a template for replication. Both gRNA and sgRNA are capped and polyadenylated.

HEK293 cells are an excellent cellular model to study SINV, as its infection exhibits all the expected molecular signatures, including: 1) active viral replication (Figure 1C and S1B-C); 2) shut off of host protein synthesis while viral proteins are massively produced (Figure 1C and S1B); 3) phosphorylation of the eukaryotic initiation factor 2 subunit alpha (EIF2α) due to the activation of the protein kinase R (PKR) (Ventoso et al., 2006) (Figure 1D); and 4) formation of cytoplasmic foci enriched in viral RNA and proteins, commonly known as viral replication factories (Figure S1C). Infection of these cells with SINV caused a strong induction of the antiviral programme, including β-interferon (β-IFN) and interferon inducible factors, which reflects the existence of active antiviral sensors and effectors (Figure S1D). Importantly, we can achieve a synchronised and near-complete infection of the cell culture with relatively low number of viral particles [multiplicity of infection (MOI)] (Figure S1E), reducing cell-to-cell variability and biological noise.

Pilot RNA-IC experiments in uninfected (mock) and SINV-infected cells revealed the isolation of a protein pool matching that previously observed for human RBPs (Baltz et al., 2012; Castello et al., 2012), which strongly differed from the total proteome (Figure 1E). No proteins were detected in non-irradiated samples, demonstrating the UV-dependency of RNA-IC. Infection did not induce major alterations in the protein pattern observed by silver staining, which correspond to the most abundant, housekeeping RBPs (Figure 1E). However, other less predominant bands displayed substantial differences, calling for indepth proteomic analysis. Oligo(dT) capture led to the isolation of both host and SINV RNAs in infected cells (Figure 1F), which is expected as gRNA and sgRNA are polyadenylated. The amount of viral RNA captured increased throughout the infection, confirming the active replication of SINV in HEK293 cells.

### SINV causes a global remodelling of the cellular RBPome

To allow accurate quantification of RBPs associated with poly(A) RNA under different physiological conditions, we developed ‘comparative RNA-IC’ by combining the original protocol (Castello et al., 2013) with stable isotope labelling by amino acids in cell culture (SILAC) (Figure 1A) (Ong and Mann, 2006). In brief, cells were grown in presence of light (normal), medium or heavy amino acids with incorporation efficiency >98%. Labelled cells were infected with SINV and irradiated with UV light at 4 and 18 hours post infection (hpi), using uninfected cells as a control (Figure 1A). These times correlate with important events in the SINV biological cycle; i.e. at 4 hpi, viral gene expression co-exists with host protein synthesis, while the proteins synthesised at 18 hpi are almost exclusively viral (Figure 1C).

SILAC labels were permutated between the three conditions (i.e. uninfected, 4 hpi and 18 hpi) in the three biological replicates to correct for possible isotope-dependent effects. After lysis, aliquots were stored for parallel transcriptomic and whole proteome analyses (see below). We combined equal amounts of the lysates from the three conditions prior to the oligo(dT) capture (Figure 1A) and eluates from the oligo(dT) capture of these mixed samples were analysed by quantitative proteomics. Protein intensity ratios between condition pairs were computed and the significance of each protein intensity change was estimated using a moderated t-test (Figure 2A-D and S2A-B). We used a previously described semi-quantitative method for the cases in which an intensity value was missing (‘zero’) in one of the two conditions leading to ‘infinite’ or ‘zero’ ratios (Sysoev et al., 2016).

**Figure 2.**
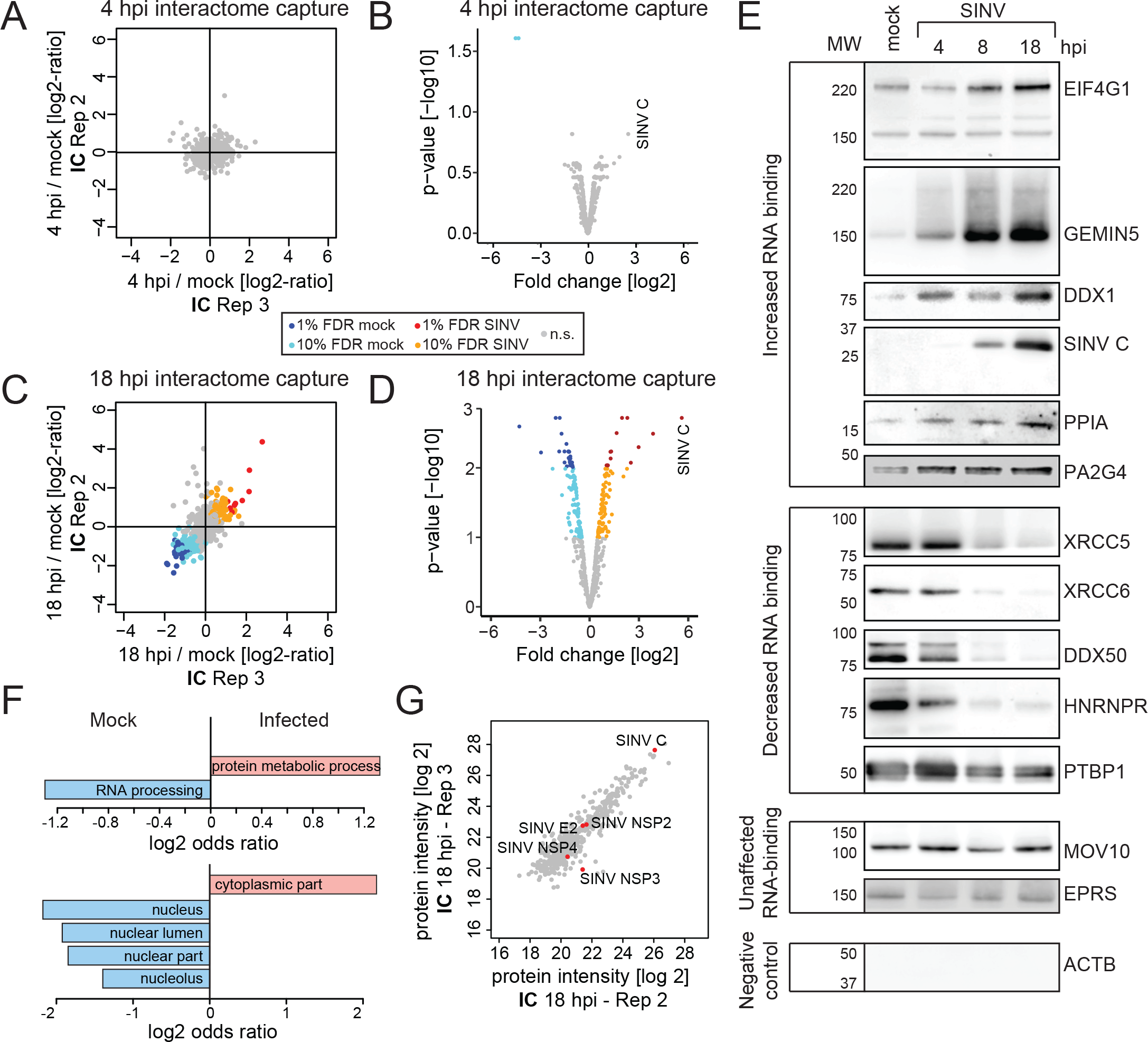
Analysis of the RNA-bound proteome in SINV-infected HEK293 cells by RNA-IC. A) Scatter plot compiling the intensity ratio between 4 hours post infection (hpi) and uninfected conditions of each protein (dots) in the eluates of two biological replicates of RNA-IC. B) Volcano plot comparing the log2 fold change of each protein between 4 hpi and uninfected conditions and the p-value of this change across the three biological replicates. C) As in (A) but comparing 18 hpi and uninfected cells. D) As in (B) but comparing 18 hpi and uninfected cells. E) Western blotting analysis with specific antibodies of the eluates of a representative RNA-IC experiment in SINV-infected HEK293 cells. F) Molecular function (upper panel) and cellular component (bottom panel) gene ontology (GO) term enrichment analysis of the RBPs stimulated (salmon) against those inhibited (blue) by SINV at 18 hpi. G) Representative scatter plot comparing the raw intensity of each protein in the eluates of two RNA-IC replicates from cells infected for 18 h. FDR, false discovery rate; n.s. non-significant.

We identified a total of 794 proteins; 91% of which were already annotated by the gene ontology term ‘RNA-binding’ or/and were previously reported to be RBPs in eukaryotic cells by RNA-IC (Hentze et al., 2018). Hence, the protein composition of our dataset largely resembles that of previously established RBPomes. Most cellular RBPs remained unaltered at 4 hpi with the exception of 17 RBPs (~2% of the identified RBPome) (Figure 2A-B, S2A and Table S1). 15 of these were detected exclusively by the semi-quantitative method due to the lack of protein intensity value in one condition, reflecting possible ‘on-off’ and ‘off-on’ states (Table S1). By contrast, SINV caused a pervasive remodelling of the RBPome at 18 hpi (Figure 2C-D and S2B), since 236 RBPs (~30%) displayed altered RNA-binding activities (48 RBPs with 1% false discovery rate [FDR], 167 with 10% FDR and 21 by the semi-quantitative analysis) (Table S1). The 245 RBPs with differential RNA-binding activity in SINV-infected cells (4 and 18 hpi) are referred to here as ‘dynamic RBPs’. Interestingly, 181 out of 245 dynamic RBPs lack classical RBDs (Castello et al., 2012; Lunde et al., 2007), suggesting that unconventional RBPs may have biological roles in infection.

To validate these results, we applied RNA-IC to cells infected with SINV but, in this case, the eluates were analysed by western blotting. We selected nine dynamic RBPs falling into three statistical categories (i.e. four with 1% FDR, four with 10% FDR and one with nonsignificant changes). As positive control, we monitored the SINV capsid (C), which is a viral RBP that accumulates throughout the infection. We also analysed two ‘non-dynamic’ RBPs (MOV10 and bifunctional glutamate/proline-tRNA ligase, EPRS) and β-actin (ACTB) as a protein lacking RNA-binding activity. The RNA-binding behaviour of each protein fully matched the proteomic outcome, including those classified with 10% FDR. The eukaryotic initiation factor (EIF) 4G1, GEMIN5, DDX1, proliferation-associated protein 2G4 (PA2G4) and peptidyl-prolyl cis-trans isomerase A (PPIA) exhibited increased interaction with RNA after infection; whereas the opposite was observed for X-ray repair cross-complementing protein 5 (XRCC5), XRCC6, DDX50 and polypyrimidine tract-binding protein 1 (PTBP1) (Figure 2E). These changes increased progressively throughout the infection. The proteomic data assigned a non-significant decrease of RNA-binding activity to HNRNPR (Table S1); however, the reduced activity of this protein was apparent by western blotting (Figure 2E), suggesting that our dataset may contain false negatives. Nonetheless, the excellent agreement between the proteomic and western blotting data supports the high quality of our results.

### Determination of the RBP networks responding to SINV infection

Among the 245 dynamic RBPs, 236 were identified at 18 hpi, 133 presented reduced and 103 increased association with RNA and they are referred to as ‘inhibited’ and ‘stimulated’ RBPs, respectively. Most of the RBPs inhibited by SINV were linked to nuclear processes such as RNA processing and export (Figure 2F and S2C). Inhibition of nuclear RNA metabolism by cytoplasmic viruses has been extensively reported although it is still poorly understood (Akhrymuk et al., 2012; Gorchakov et al., 2005; Lloyd, 2015), and the inhibition of cellular RBPs is likely contributing to this phenomenon. Conversely, most stimulated RBPs are cytoplasmic and are linked to protein synthesis, 5’ to 3’ RNA degradation, RNA transport, protein metabolism and antiviral response (Figure 2F and S2D). Nonetheless, several nuclear RBPs, including splicing factors, are stimulated upon infection, indicating that the above statements have notable exceptions.

Interestingly, several RBPs involved in translation were stimulated at 18 hpi despite the shut off of host protein synthesis (Figure 1C and S1B). These include eukaryotic initiation factors (EIF4G1, EIF4G2, EIF4G3, EIF3C, EIF3D, EIF3E, EIF3G, EIF3J and EIF5B), elongation factors (EEF1G, EEF2, EEF1A1) and ribosomal proteins (RPS27, RPS25, RPS15A, RPS12, RPL36, RPL28, RPL10, RPL29, RPL39, RPL8, RPL4, RPL9 and RPL14). This enhancement is likely due to the high translational activity of SINV RNAs (Figure 1C) (Frolov and Schlesinger, 1996). Conversely, four ribosomal proteins (RPS3, RPS10, RPS28 and RPL7L1) showed reduced RNA-binding activity (Table S1). The essential components of the cap-dependent translation initiation machinery, EIF4A1 and EIF4E, were not stimulated by the infection in spite of the activation of their protein partner EIF4G1 (Table S1). This agrees with previous data showing that these two initiation factors do not participate in SINV sgRNA translation (Carrasco et al., 2018; Castello et al., 2006; Garcia-Moreno et al., 2015; Garcia-Moreno et al., 2013). A recent report showed that EIF3D is a cap-binding protein that substitutes EIF4F in the translation of specific mRNA pools (Lee et al., 2016). EIF3D is stimulated by SINV, and thus its potential contribution to SINV RNA translation deserves further consideration. Moreover, 87 dynamic RBPs were recently reported to associate with ribosomes in a proteomic analysis using Flag-tagged ribosomal proteins in mouse cells (Table S2) (Simsek et al., 2017). The existence of ‘specialised ribosomes’ in infected cells has been proposed; however, experimental evidence is currently sparse (Au and Jan, 2014). Our results indicate that the composition of ribosomes and the scope of proteins associated with them may strongly differ between infected and uninfected cells, possibly resulting in differential translational properties.

RNA-IC uncovered 16 dynamic RNA helicases (Table S2); 13 of which were inhibited upon infection. RNA helicases are fundamental at virtually every stage of RNA metabolism, since they mediate the remodelling of RNPs (Chen and Shyu, 2014). Hence, it is expected that their inhibition will have important consequences in RNA metabolism. Only 3 helicases were stimulated by SINV (DDX1, DHX57 and DHX29) (Figure 2E and Table S2). DDX1 is part of the tRNA ligase complex (Popow et al., 2011) and interestingly the ligase subunit, RTCB, is also stimulated by SINV (Table S1), as will be discussed below. DHX29 enhances 48S complex formation on SINV sgRNA in reconstituted *in vitro* systems (Skabkin et al., 2010), and the stimulation of this helicase reported by RNA-IC supports its potential contribution to viral translation in infected cells.

Notably, a defined subset of antiviral factors displayed increased RNA-binding activity in response to SINV infection, suggesting activation through PAMP recognition. These RBPs include the interferon-inducible protein 16 (IFI16), interferon-induced protein with tetratricopeptide repeats 5 (IFIT5), E3 ubiquitin/ISG15 ligase TRIM25, TRIM56 and zinc finger CCCH-type antiviral protein 1 (ZC3HAV1 or ZAP) (Table S1). IFI16 was previously described to bind double stranded DNA in cells infected with DNA viruses (Ni et al., 2016). Our data reveal that IFI16 also binds to RNA, and it is activated early after SINV infection (4 hpi). This agrees with the recently described ability of IFI16 to restrict RNA virus infection (Thompson et al., 2014). These findings highlight the capacity of RNA-IC to identify antiviral factors responding to a given RNA virus.

Interestingly, RNA-IC also identified viral RBPs associated with poly(A) RNA in infected cells. These include the known viral RBPs (i.e. RNA helicase NSP2, the RNA polymerase NSP4 and the capsid protein) and, unexpectedly, also NSP3 and E2 (Figure 2G and S2E). NSP3 was only quantified in two replicates (Figure S2E), and thus its interaction with RNA requires experimental confirmation. The identification of E2 in RNA-IC eluates was unexpected. However, the structure of E1, E2 and C from the related VEEV provides some insights into this finding. E2 interacts with C nearby cavities that communicate with the inner part of the viral particle, where the gRNA density resides (Zhang et al., 2011). It is thus plausible that E2 interacts transitorily or stochastically with the viral RNA in the context of the highly packed capsid.

### RBP responses to SINV are not caused by changes in protein abundance

Changes detected by RNA-IC can be a consequence of matching alterations in protein abundance (Sysoev et al., 2016). To assess this possibility globally, we analysed the total proteome (inputs of the RNA-IC experiments, Figure 1A) by quantitative proteomics (Figure 3). Importantly, SINV infection did not cause noticeable alterations in host RBP levels, even at 18 hpi (Figure 3A-C, S3A-C and Table S3). In agreement, coomassie staining did not show major protein fluctuations in uninfected versus infected cells except for SINV C (Figure 3D). The lack of changes in protein levels, even for dynamic RBPs, was confirmed by western blotting (Figure 3E). Taken together, these data confirm that the variations in RNA-binding activity of cellular RBPs are not globally caused by alterations in the host protein abundance.

**Figure 3.**
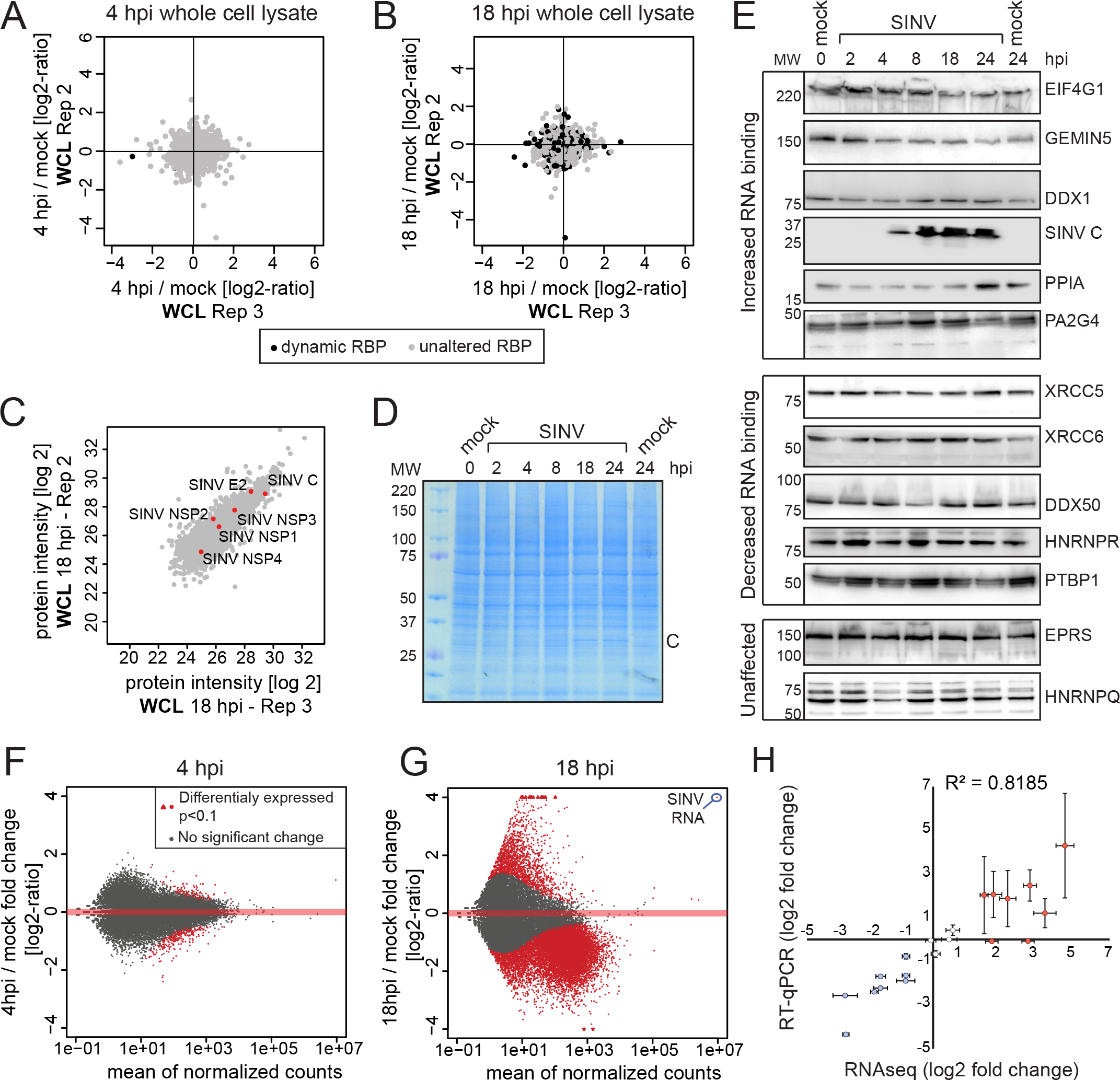
Proteomic and transcriptomic analyses of whole SINV-infected cell lysates. A) Scatter plot compiling the intensity ratio between 4 hpi and uninfected conditions of each protein (dots) in the inputs (total proteome) of two biological replicates of RNA-IC (scatter plots of the three biological replicates in Figure S3). Black dots represent proteins significantly enriched in either 4 hpi or uninfected conditions in the RNA-IC experiment (Figure 2 and Table S1). B) As in (A) but between 18 hpi and uninfected conditions. C) Scatter plot comparing the intensity of each protein in the inputs (total proteome) of two RNA-IC replicates from cells infected for 18 h (scatter plots of the three biological replicates in Figure S3). D) Representative Coomassie blue staining of the whole cell lysate of cells infected with SINV for 2, 4, 8, 18 or 24 h. E) Western blotting analysis with specific antibodies of the whole cell lysates (inputs) from SINV-infected HEK293 cells. F) MA plot comparing the read coverage and the log2 fold change between 4 hpi and uninfected cells of each transcript detected in the RNAseq experiment. Red dots represent RNAs enriched in one of the two conditions with p<0.1. G) As in (F) but between 18 hpi and uninfected cells. H) Correlation of the RNAseq and RT-qPCR data by plotting the log2 fold change for randomly selected transcripts by the two methods. Error bars represent standard error of three independent experiments.

### The transcriptome undergoes pervasive alterations in SINV-infected cells

Mechanistically, the activity of host RBPs can also be altered by changes in the availability of their target RNAs. To profile the transcriptome of uninfected versus SINV-infected cells, we performed RNA sequencing analysis of total RNA isolated from the RNA-IC input samples (Figure 1A). 4 h of SINV infection had a relatively minor impact on the host transcriptome, with only 67 and 177 up- and down-regulated RNAs, respectively (Figure 3F). By contrast, profound changes in the cellular transcriptome were observed at 18 hpi, with 12,372 differentially expressed RNAs (p<0.1; Figure 3G and S3E-F). While 10,924 RNAs had significantly lower expression in infected cells, only 1,448 RNAs were upregulated (Table S4). Hence, SINV infection causes a massive loss of cellular RNAs. Upregulated RNAs were enriched in genes annotated by the GO term ‘antiviral response’, reflecting the activation of the host defences (Figure S1D, S3E and Table S4). To validate these results by an orthogonal approach, we used RT-qPCR focusing on 20 mRNAs randomly chosen across the whole variation range. Importantly, data obtained with both techniques strongly correlated (R^2^= 0.82) (Figure 3H). In summary, availability of cellular RNA is globally reduced upon infection, correlating with the emergence of viral RNA (Figure 3G and Table S4). Decreased availability of cellular RNA is expected to contribute to the inhibition of RNA-binding activity observed for 133 RBPs (Figure 2C-D and Table S1).

### SINV infection induces subcellular redistribution of cellular RBPs

While depletion of cellular RNA can explain the reduced activity of RBPs, it cannot account for the stimulation of 103 proteins. Importantly, SINV RNAs are amongst the most abundant RNA species in the cell at 18 hpi (Figure 3G). Such abundant RNA substrate is likely to influence cellular RBPs, potentially driving the remodelling of the RBPome. SINV produces two overlapping mRNAs, gRNA and sgRNA (Figure 1B and S1A) and, consequently, the read coverage was substantially higher in the last third of the gRNA, where both transcripts overlap (Figure 4A). Both sgRNA and gRNA are substantially more abundant than the negative strand (Figure 4A). As the copy number of the negative strand is low and it lacks poly(A) tail, it should not contribute to the RNA-IC results.

**Figure 4.**
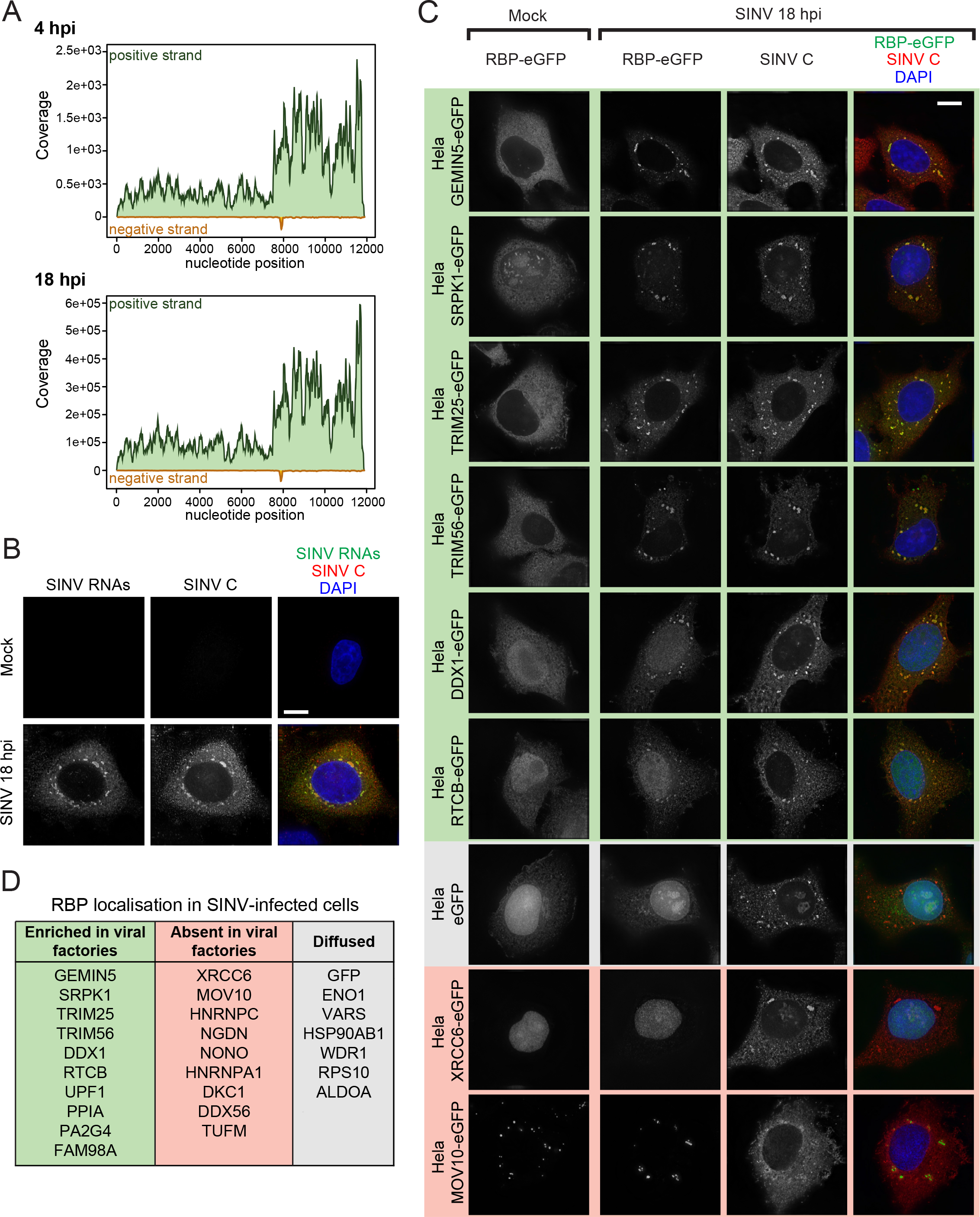
Host RBP localisation in SINV infected cells. A) Read coverage of the positive and negative RNA strand of SINV in the RNAseq analysis at 4 and 18 hpi. Note that the Y axis in both plots have different scales to facilitate the visualisation of the data. B) Localisation analysis of SINV RNA and C protein in infected HeLa cells at 18 hpi by combined *in situ* hybridisation and immunofluorescence. C) Localisation by immunofluorescence of the eGFP-fused RBPs and SINV C in HeLa cells uninfected or infected with SINV for 18 hpi. D) Summary of the observed localisation of the 26 proteins tested in panel C and Figure S4B. Scale-bar represents 10 μm.

SINV RNAs are known to accumulate in cytoplasmic compartments, so called viral replication factories, derived from the membranes of cellular organelles (Figure 4B) (Romero-Brey and Bartenschlager, 2014). These viral RNA-containing foci can explain the punctuate accumulations of poly(A) RNA in the cytosol of infected cells (Figure S4A). We thus reasoned that RBPs interacting with SINV RNAs should accumulate in these compartments. To explore this idea systematically, we generated 26 tetracycline-inducible cell lines expressing host RBPs fused to eGFP. This cell library included 16 lines expressing ‘stimulated RBPs’ and 8 expressing ‘inhibited RBPs’. A non-dynamic RBP, MOV10 (Figure 2E), and unfused eGFP were used as controls. Strikingly, 9 out of the 16 stimulated RBPs (56%) accumulated at viral replication factories demarcated by SINV C, including cytoplasmic and nuclear RBPs (Figure 4C-D and S4B). As SINV C is a good proxy for the localisation of viral RNAs (Figure 4B), RBPs retained at these foci are likely to interact with viral RNA. To our knowledge, this is the first time that the idea of viral RNA hijacking cellular RBPs in viral replication factories is assessed in a systematic way. Five additional stimulated RBPs (29%) showed diffuse localisation in cytoplasm, but were also present at the SINV C-containing foci (Figure S4B). Amongst the stimulated RBPs, only neuroguidin (NGDN), heterogeneous ribonucleoprotein A1 (HNRNPA1) and the mitochondrial Tu translation elongation factor (TUFM) (3 out of 16; 17%) were absent in the viral factories. The absence of these three proteins in the replication factories suggests that the stimulation of their RNA-binding activity may involve exclusively host RNA. Nevertheless, HNRNPA1 was shown to bind SINV RNA (LaPointe et al., 2018; Lin et al., 2009), although in our analysis it displays strictly nuclear localisation (Figure S4B). We cannot rule out that a small pool of HNRNPA1 is present in the viral factories at undetectable levels or, alternatively, that the eGFP tag is affecting HNRNPA1 localisation.

In contrast to stimulated RBPs, only one (out of 8; 12.5%) virus-inhibited RBP was enriched in the viral factories (Figure 4D and S4B). This protein, called regulator of nonsense transcripts 1 (UPF1), is a helicase involved in the non-sense mediated decay (NMD) pathway and is known to inhibit infection of alphaviruses (Balistreri et al., 2014). Conversely, 5 out of 8 (62.5%) virus-inhibited RBPs are nuclear and remained nuclear after infection (Figure 4C-D and S4B). Several of the nuclear RBPs (e.g. RPS10, DKC1, NGDN or HNRNPA1) display different degrees of delocalisation in infected cells, suggesting either release from the nuclear compartments where they reside or partial disassembly of these structures.

MOV10 is a helicase unaffected by SINV infection (Figure 2E) that localises in p-bodies (Goodier et al., 2012). To our surprise, MOV10-containing p-bodies were surrounded by viral replication factories in SINV-infected cells, in a non-overlapping fashion (Figure 4C). Our results indicate that p-bodies may crosstalk with viral replication factories.

In summary, global alterations of RBP localisation and differential availability of RNA substrates are likely jointly driving the changes in RNA-binding activity in SINV-infected cells.

### Activation of XRN1 and RNA degradation in SINV-infected cells

The loss of cellular mRNAs can be an important driving force at shaping the RBPome in SINV-infected cells (Figure 3G). However, it is unclear how the transcriptome remodelling is originated and whether it benefits viral infection. Alterations in RNA levels can be a consequence of reduced transcription and/or increased RNA degradation (Mukherjee et al., 2017). To explore which of these pathways contribute the most to RNA loss in SINV infected cells, we compared the fold change of each mRNA in our dataset to available data compiling the speed of synthesis, processing and degradation of each individual transcript (Mukherjee et al., 2017) (Figure 3G and Table S4). Transcription could explain most of the differences between uninfected and 4 hpi (Figure 5A and S5A). However, RNA degradation accounted for more than 50% of the explained variance at 18 hpi. We reasoned that this phenomenon can be a combined effect of the activation of the 5’ to 3’ RNA degradation machinery, as the exonuclease XRN1 and its interactor PAT1 homolog 1 (PATL1) are stimulated at 18 hpi (Figure S2D and Table S1), and a reduced transcriptional activity (Gorchakov et al., 2005). The role of XRN1 in virus infection is currently controversial. Some studies reported that XRN1 restricts viral replication by erasing viral RNA (Molleston and Cherry, 2017), whereas others proposed that RNA fragments resistant to XRN1 degradation can interfere with the antiviral response (Manokaran et al., 2015). In SINV-infected cells, XRN1-containing foci (p-bodies) are juxtaposed to the viral replication factories (Figure 5B), as observed for MOV10 (Figure 4C). To assess XRN1 effects in SINV infection, we generated XRN1 knock out cell lines and studied the fitness of the chimeric SINV-mCherry (Figure 1B). To our surprise, XRN1 knock out cells were completely refractory to SINV-mCherry infection, while partial knock out led to an intermediate phenotype (Figure 5C). The fact that SINV fails to infect cells lacking XRN1 suggests that its activity is critical for the infection. Importantly, the identification of XRN1 as an essential host RBP for SINV highlights the capacity of RNA-IC to uncover in a global scale host factors implicated in virus infection.

**Figure 5.**
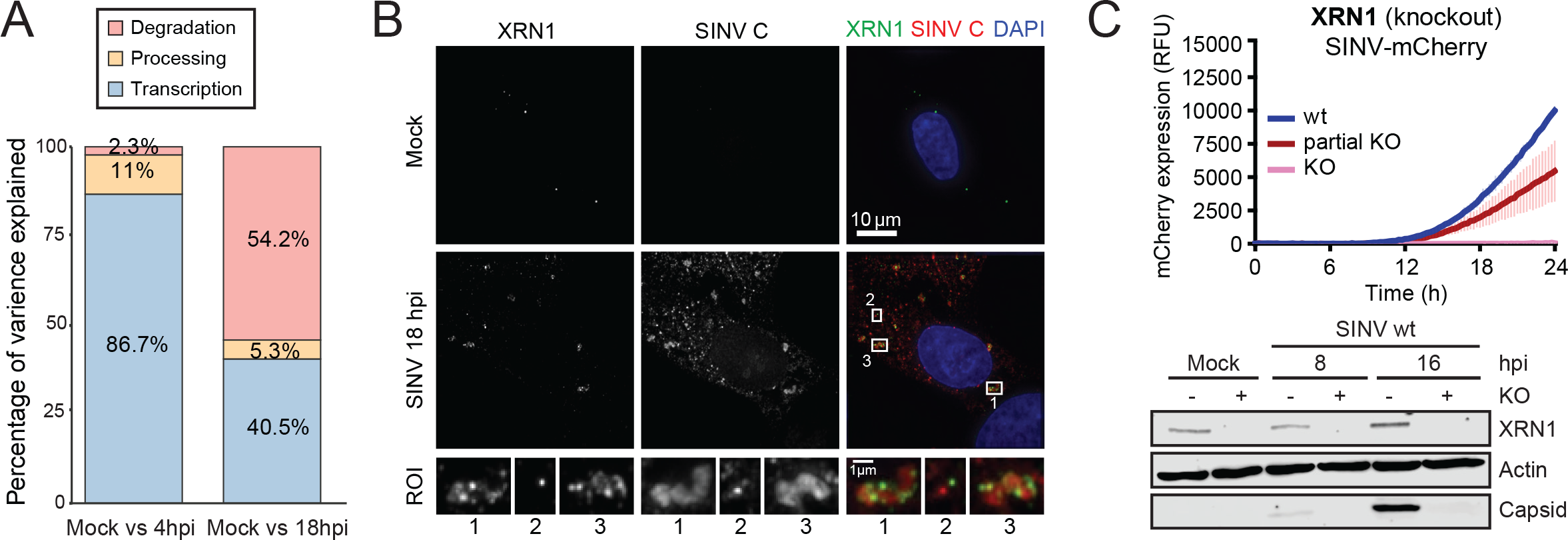
The exonuclease XRN1 in cells infected with SINV. A) Analysis of the contribution of transcription, processing and degradation to the transcriptome of cells infected with SINV. The RNA expression data (Figure 3F-G and Table S4) was compared with the speed of RNA synthesis, processing and degradation independently measured for each transcript in (Mukherjee et al., 2017). ANOVA was used to estimate the percentage of variance explained by each RNA biological process (i.e. transcription, processing and degradation) in the different conditions. B) Immunolocalisation of XRN1 and SINV C using specific antibodies. ROI, region of interest. C) Upper panel shows the fluorescence derived from mCherry in XRN1 knock out and control HEK293 cells infected with SINV-mCherry. Red fluorescence was measured every 15 min in a plate reader with atmospheric control (5% CO2 and 37°C). RFU, relative fluorescence units. Bottom panel shows the expression of SINV C in XRN1 knock out and wild type HEK293 cells analysed by western blotting.

### RBPome responses are biologically important

To determine in a broader extent whether RBP responses are functionally important, we sought to study the impact of dynamic RBPs on virus infection. The ligase RTCB, together with DDX1, FAM98A, FAM98B, CGI-99 and Ashwin (ASHWN) form the tRNA ligase complex (TRLC). Two components of this complex, RTCB and DDX1, displayed enhanced RNA-binding activity (Table S1). Moreover, these two proteins and the nuclear factor FAM98A accumulated in the viral replication factories (Figure 4C and S4B). TRLC ligates the exons during the splicing of tRNAs (Popow et al., 2011; Popow et al., 2014) and a recent study showed that it also mediates the cytoplasmic splicing of *xbp1* mRNA upon unfolded protein response (Jurkin et al., 2014). RTCB mediates the unusual ligation of 3´-phospate or 2’, 3´-cyclic phosphate to a 5´-hydroxyl end, which are generated by a limited repertoire of cellular endonucleases that includes the serine/threonine-protein kinase/endoribonuclease IRE1 (IRE1α) (Jurkin et al., 2014; Popow et al., 2011; Popow et al., 2014). SINV has been proposed to cause unfolded protein response (Rathore et al., 2013), which is compatible with the activation of IRE1α and TRLC in infected cells. To assess the relevance of this post-transcriptional circuit in SINV-mCherry-infected cells, we used a specific inhibitor (4μ8C) against the endonuclease IREI±. Notably, this compound strongly reduced viral fitness in low, non-cytotoxic concentrations (Figure 6A and S6A). Hence, our data suggest that IRE1± and TRLC are positively contributing to SINV infection, although the precise mechanism by which this occurs should be further investigated.

**Figure 6.**
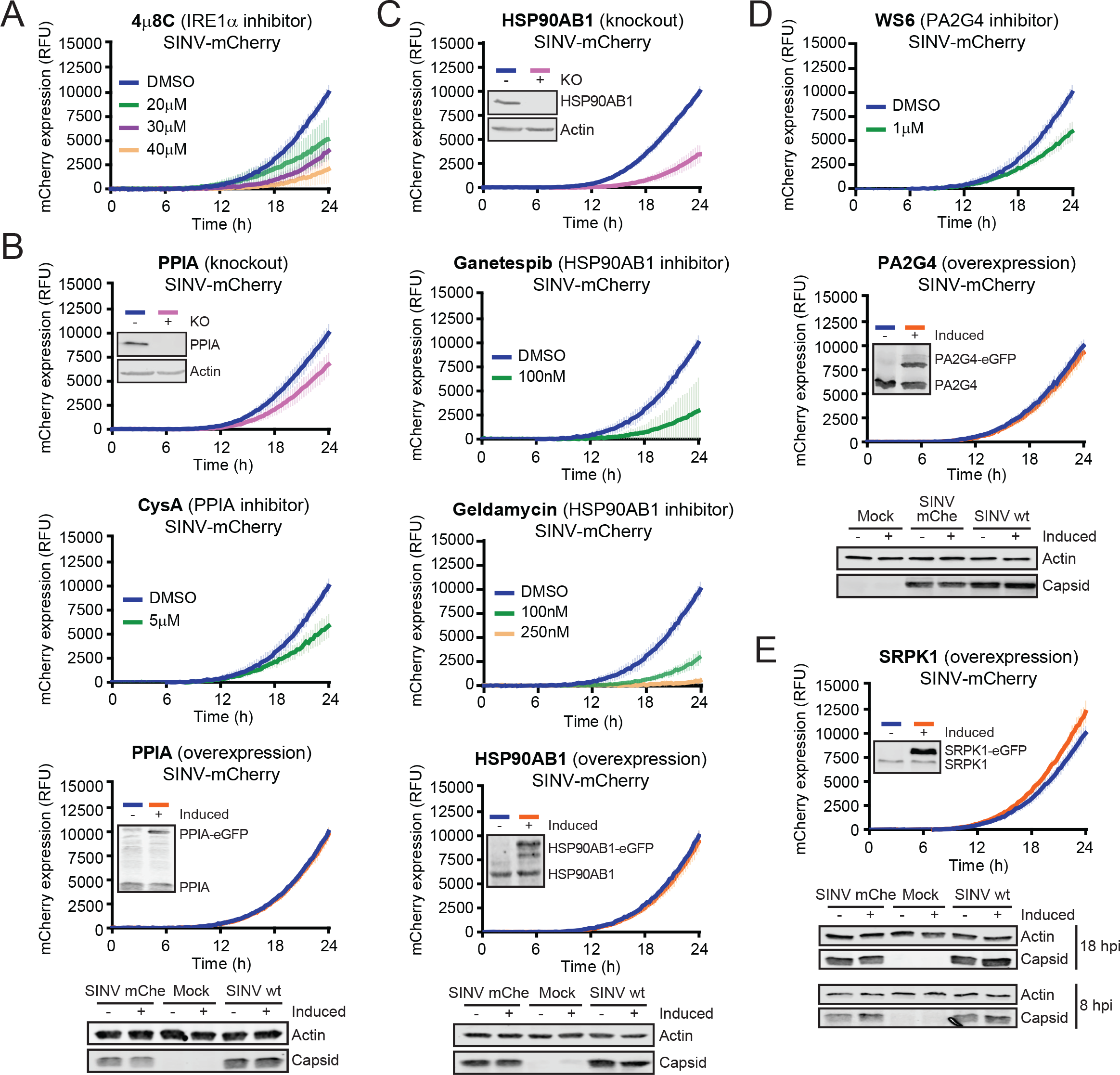
Impact of stimulated-RBPs in SINV infection. A) Expression of mCherry in HEK293 cells infected with SINV-mCherry and treated or not with the IREI± inhibitor, 4μ8C. Red fluorescence was measured as in Figure 5C. B) As in (A) but with PPIA knock out cells (first panel), the PPIA inhibitor cyclosporine A (CysA) (second panel) and cells overexpressing PPIA-eGFP (third panel). Knock out and overexpression of PPIA was assessed by Western blotting with specific antibodies. Bottom panel shows SINV C accumulation in cells overexpressing PPIA and infected with either SINV or SINV-mCherry, analysed by Western blotting. C) mCherry expression in HSP90AB1 knock out cells (first panel), in cells treated with inhibitors of HSP90 (ganetespib and geldamycin, second and third panels) or in cells overexpressing HSP90AB1-eGFP (fourth panel) and infected with SINV-mCherry. Red fluorescence was measured as in Figure 5C. Knock out and overexpression of HSP90AB1 was assessed by Western blotting with specific antibodies. Bottom panel shows SINV C accumulation in cells overexpressing HSP90AB1 and infected with either SINV or SINV-mCherry, analysed by Western blotting. D) As in (A) but the PA2G4 inhibitor WS6 (first panel) and cells overexpressing PA2G4-eGFP (second panel). Third panel shows SINV C accumulation in cells overexpressing PA2G4 and infected with either SINV or SINV-mcherry, analysed by Western blotting. E) As in (A) but with cells overexpressing SRPK1 (first panel). Overexpression of SRPK1 was assessed by Western blotting with specific antibodies. Second panel shows SINV C accumulation in cells overexpressing SRPK1 and infected with either SINV or SINV-mcherry for 8 and 18 h, analysed by Western blotting. Dox, doxycycline, which induces the (over)expression of the RBP-eGFP fusion protein in HEK293 Flp-In TREx cells. SINV-mChe, SINV-mCherry.

The peptidyl-prolyl cis-trans isomerase A (PPIA, also called cyclophilin A) has also been classified as an RBP by RNA-IC studies, together with other six members of the PPI family (Hentze et al., 2018). These enzymes switch proline conformation and modulate protein activity, with important regulatory consequences for RNA metabolism (Mesa et al., 2008). PPIA is essential for hepatitis C virus as it modulates NS5A activity during replication. Hence, the inhibition of PPIA has been exploited as a host-based antiviral therapy (Rupp and Bartenschlager, 2014). PPIA is also important for the infection of human immunodeficiency virus (HIV), influenza virus A and hepatitis B virus (Li et al., 2007; Liu et al., 2016; Zhou et al., 2012). RNA-IC revealed that PPIA is the sole PPI enzyme that responds to SINV infection (Table S1). In addition, PPIA is recruited to the viral replication factories (Figure S4B). Interestingly, SINV-mCherry infection is delayed by both the ablation of PPIA and the treatment with the PPIA inhibitor cyclosporine A at non-toxic concentrations (Figure 6B and S6A). However, PPIA-eGFP overexpression had no effect in SINV-mCherry fitness (Figure 6C, bottom panel).

The interaction of heat shock protein 90AB1 (HSP90AB1) with RNA is enhanced by SINV infection (Table S1). This protein has been identified as an RBP in numerous RNA-IC studies (Hentze et al., 2018) and its RBD has been located in a discrete region at its C-terminal domain (Figure S6B) (Castello et al., 2016). Chaperones from the HSP90 family play important roles in the remodelling of RNPs and have been linked to virus infection (Geller et al., 2012; Iwasaki et al., 2010). To address whether HSP90AB1 is involved in SINV infection, we generated a knockout cell line. Notably, the kinetics of SINV-mCherry infection was substantially delayed in cells lacking HSP90AB1 (Figure 6C, upper panel), even though homologues of this protein exist (i.e. HSP90AA1, HSP90AA2, HSP90B1 and TRAP1). The ability of HSP90AB1 to promote SINV infection was confirmed using the specific inhibitors ganetespib and geldanamycin. Treatment with non-toxic concentrations of these compounds abrogated SINV-mCherry infection (Figure 6C, middle panels, and S6A). However, HSP90AB1 overexpression had no effect in SINV-mCherry fitness (Figure 6C, bottom panel). HSP90AB1 is abundant in the cell and thus overexpression may have limited effects. The implication of PPIA and HSP90 in the biological cycle of a variety of unrelated viruses highlights these proteins as master regulators of infection, representing potential targets for broad-spectrum antivirals (Rupp and Bartenschlager, 2014; Wang et al., 2017). Proliferation-associated protein 2G4 (PA2G4) RNA-binding activity was also enhanced by SINV (Table S1). This protein associates with ribosomes (Table S2) (Simsek et al., 2017) and regulates the cap-independent translation of footh-and-mouth disease virus (FMDV) RNA (Monie et al., 2007). Treatment with the PA2G4-specific inhibitor WS6 hampered SINV-mCherry fitness (Figure 6D, upper panel), suggesting that this protein promotes SINV infection. Overexpression did not cause any effect as with previous examples (Figure 6D, bottom panel). The possibility that PA2G4 contributes to the non-canonical, cap-dependent translation of SINV RNAs should be further investigated.

SRSF Protein Kinase 1 (SRPK1) phosphorylates the RS repeats present in SR proteins, which are involved in alternative splicing regulation, RNA export and stability (Howard and Sanford, 2015). SINV infection stimulates the RNA-binding activity of SRPK1 (Table S1). We hypothesise that the recruitment of this protein to the viral replication factories (Figure 4C) might i) act as a sponging mechanism, preventing it from acting elsewhere or ii) promote phosphorylation of proteins in the context of the viral replication factories. An earlier study suggested that inhibition of SRPK1 hampers SINV and HIV infection (Fukuhara et al., 2006). To complement these experiments and distinguish between the proposed mechanisms of action, we overexpressed SRPK1-eGFP reasoning that by increasing the levels of this protein we would compromise the ‘sponge’ mechanism. Notably, SRPK1-eGFP overexpression enhanced SINV fitness (Figure 6E), suggesting that the ‘sponge’ mechanism is improbable and that it is more plausible that SRPK1 is actively contributing to viral replication in the viral factories. Future work should determine the molecular targets of SRPK1 in the viral replication factories, and whether its kinase activity also contributes to the differential RNA-binding activity of dynamic RBPs.

We tested the effects of the overexpression of nine additional responsive or inhibited RBPs fused to eGFP (Figure S6C and D). Phenotypes in viral fitness ranged from non-existent (ALDOA, XRCC6, RPS10, MOV10 and NGDN), to mild (RPS27, NONO and DKC1). The lack of phenotypic effects in overexpression experiments does not discard that the protein actually participates in SINV infection, as shown before with HSP90AB1, PPIA and PA2G4. Nevertheless, RBPs for which overexpression caused mild effects in infection have potential for future in detail analyses.

The family of tripartite motif-containing (TRIM) proteins comprises more than 75 members endowed with E3 ubiquitin ligase activity that are known to modulate the host antiviral response (Versteeg et al., 2013). RNA-IC studies have discovered RNA-binding activities in few TRIM proteins, including TRIM25, TRIM56, TRIM28 and TRIM71 (Hentze et al., 2018). Notably, SINV infection enhanced TRIM25 interaction with RNA (Table S1), correlating with its relocalisation to the viral replication factories (Figure 4C). TRIM25 was proposed to interact with dengue virus RNA (Manokaran et al., 2015); however, this analysis employed native immunoprecipitation (IP) that cannot distinguish between direct and indirect protein-RNA interactions (e.g. through protein-protein interactions). To prove experimentally that TRIM25 interacts directly with SINV RNA, cells expressing TRIM25-eGFP were infected with SINV and irradiated with UV light to induce protein-RNA crosslinks. TRIM25-eGFP was IP under very stringent conditions (500 mM NaCl and 0.1% SDS) and co-isolated RNA was analysed by RT-PCR using specific primers against SINV RNA. A band matching the expected PCR product was amplified from TRIM25-eGFP IPs only in infected cells (Figure 7A), confirming its interaction with viral RNA. No PCR product was observed in the negative controls, i.e. unfused eGFP or XRCC6-eGFP. TRIM25 RNA-binding activity is mediated by a short peptide present in its PRY domain and interaction with RNA enhances its E3 ubiquitin ligase activity (Choudhury et al., 2017). Thus, the enhanced RNA-binding activity of TRIM25 in SINV-infected cells is expected to increase its E3 ubiquitin ligase activity, which is known to stimulate key effectors of the antiviral response such as RIGI and ZC3HAV1 (Gack et al., 2007; Li et al., 2017; Zheng et al., 2017). Hence, RBP ubiquitination may also affect the RNA-binding activity of dynamic RBPs in the context of the viral factory. Another SINV-stimulated RBP is TRIM56, which is known to bind double stranded DNA. However, it enhances the antiviral response not only in cells infected with DNA viruses but also with RNA viruses (Seo et al., 2018; Shen et al., 2012; Tsuchida et al., 2010; Versteeg et al., 2013). RNA-IC thus complements these results revealing that TRIM56 interacts directly with RNA in SINV-infected cells (Table S1). Moreover, TRIM56 also re-localises to the viral replication factories (Figure 4C). Strikingly, overexpression of both TRIM25-eGFP and TRIM56-eGFP strongly reduced SINV fitness (Figure 7B), confirming that these RBPs are part of the cell’s molecular arsenal to restrict virus infection. 160 out of the 245 SINV-responsive RBPs lack previous connections to virus infection, based on our recent analysis of protein annotation and automated literature mining (Garcia-Moreno et al., 2018). Hence, our dataset likely contains numerous pro- and antiviral RBPs to be discovered.

**Figure 7.**
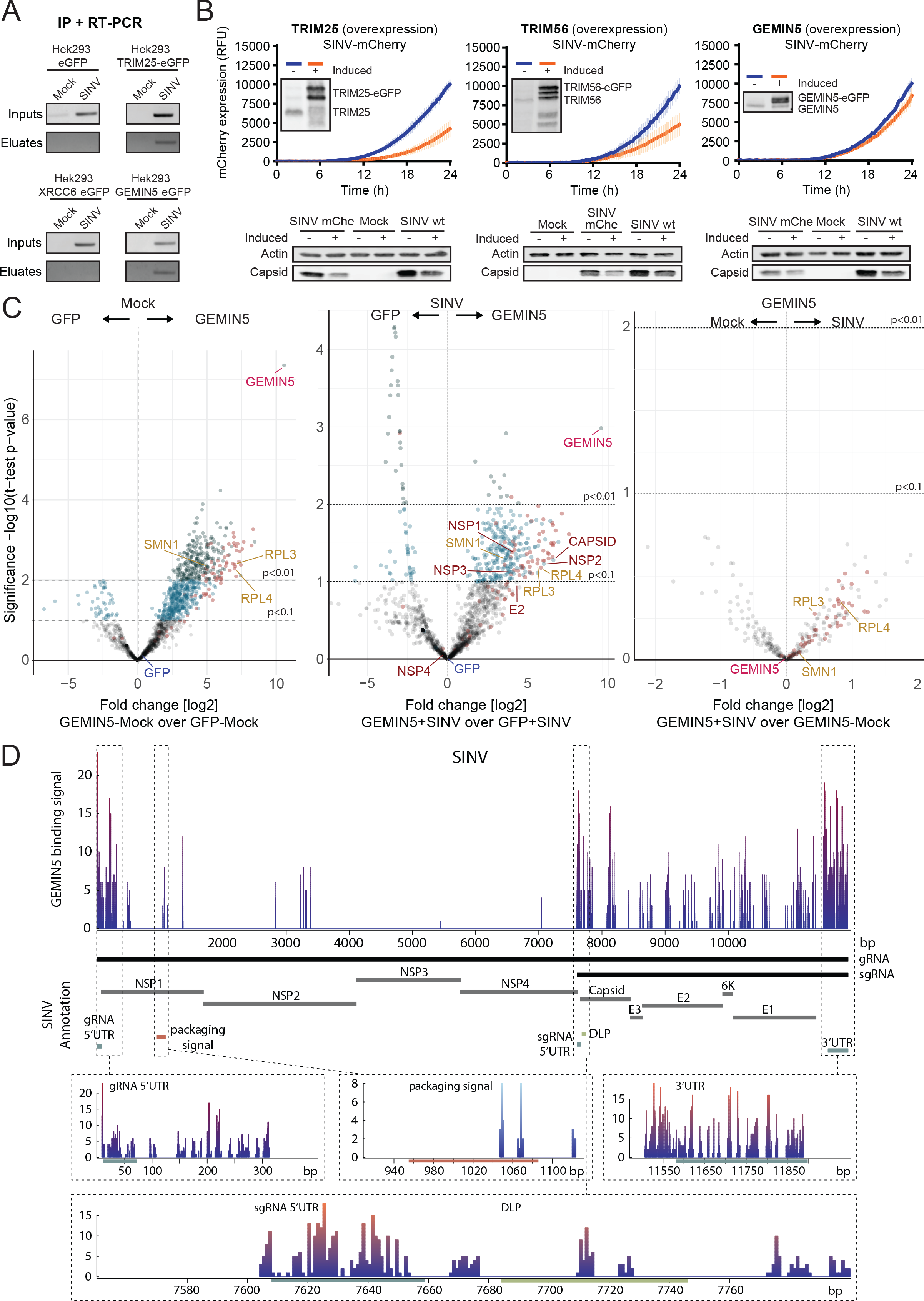
Effects of RBPs with antiviral potential in SINV infection. A) UV crosslinking and immunoprecipitation of TRIM25-eGFP, GEMIN5-eGFP, XRCC6-eGFP or unfused eGFP in cells infected or not with SINV. The presence SINV RNA in eluates and inputs was detected by RT-PCR using specific primers against SINV RNAs (overlapping sequence). B) Relative mCherry fluorescence produced in cells overexpressing TRIM25-eGFP (left upper panel), TRIM56-eGFP (middle upper panel), GEMIN5-eGFP (upper right panel) and infected with SINV-mCherry. Red fluorescence was measured as in Figure 5C. Overexpression of these proteins was assessed by Western blotting with specific antibodies. Bottom panel shows SINV C accumulation in cells overexpressing these proteins and infected with either SINV or SINV-mcherry. SINV C was analysed by Western blotting using β-actin (ACTB) as loading control. C) Volcano plots comparing the intensity of proteins in GEMIN5-eGFP vs unfused eGFP IP in uninfected (left panel) and infected cells (middle panel); every dot represents a protein. Dark green dots are proteins enriched in either GEMIN5-eGFP or unfused eGFP with p<0.01; blue dots are those enriched with p<0.1 and grey dots represent non-enriched proteins. Pink dots represent ribosomal proteins. Right panel shows a volcano plot comparing the intensity of proteins in GEMIN5 IPs in infected vs uninfected cells. D) iCLIP analysis of GEMIN5 binding sites on SINV RNA. The upper panel shows the reads mapping to SINV positive-stranded RNA. Discontinuous boxes below the plot delimit relevant regulatory regions within the SINV RNA. Black boxes demark the position of gRNA and sgRNA, grey boxes demark the position of viral proteins and coloured boxes delimit the position of relevant regulatory regions. The reads mapping to the 5’ and 3’ regions of the gRNA and sgRNA (including the downstream loop or DLP) as well as the C-binding site (packaging signal) are zoomed in in bottom panels.

### GEMIN5 binds to key regulatory sequences in viral RNAs and regulates SINV protein expression

GEMIN5 is a member of the survival motor neuron (SMN) complex, which mediates the assembly of the small nuclear (sn)RNPs (Gubitz et al., 2002). The RNA-binding activity of this protein is strongly stimulated by SINV infection and it redistributed to the viral replication factories (Figure 2E and 4C). To our surprise, neither the other GEMIN proteins nor the SMN proteins, known molecular partners of GEMIN5, were stimulated by SINV (Table S1), suggesting a GEMIN5-specific response. This agrees with the earlier discovery of a free pool of GEMIN5 (Battle et al., 2007). Further work reported that GEMIN5 is a global regulator of protein synthesis and is cleaved upon FMDV infection (Francisco-Velilla et al., 2016; Pacheco et al., 2009; Pineiro et al., 2015). Overexpression of GEMIN5-eGFP caused a slight but highly reproducible delay of mCherry production and strongly inhibited capsid synthesis (Figure 7B). GEMIN5 impacts protein synthesis at the translation elongation step through its direct interaction with the 60S ribosomal subunit and, in particular, with RPL3 and RPL4 (Francisco-Velilla et al., 2016). In agreement, GEMIN5 co-precipitates with purified ribosomes (Table S2) (Simsek et al., 2017). Protein-protein interaction analysis of GEMIN5-eGFP followed by proteomics confirmed that, in our experimental settings, it interacts with RPL3 and RPL4 as well as with other components of the ribosome, especially from the 60S subunit (Figure 7C, pink dots in left panel, S7A-B and Table S5). Moreover, the interaction with the ribosomes is maintained in SINV-infected cells (Figure 7C, pink dots in middle and right panels). We noticed that GEMIN5 is by far the most enriched protein in the IPs and that its iBAQ score is significantly higher than that of eGFP, suggesting that GEMIN5-eGFP interacts with the endogenous GEMIN5 likely forming oligomers through its C-terminus coiled-coil structure, as previously proposed (Xu et al., 2016). Moreover, our data showed that GEMIN5 may interact with various viral proteins, chiefly with NSP1, NSP2, NSP3 and SINV C (Figure 7C, middle panel). The biological implications of these interactions in the modulation of the RNA-binding activity of GEMIN5 deserve future considerations.

To test whether GEMIN5 binds SINV RNA directly, we performed IP and RT-PCR analysis as outlined above. A PCR product of the expected size was amplified in GEMIN5-eGFP eluates, while the negative controls were remarkably clean (Figure 7A). To determine the exact positions within the viral RNA bound by GEMIN5, we employed single-nucleotide resolution crosslinking and immunoprecipitation (iCLIP) followed by sequencing (König et al., 2010). The binding peaks with highest coverage mapped to the 5’ region of the gRNA and sgRNA as well as to the 3’ untranslated region (UTR), which is shared by the two viral RNAs (Figure 7D). Indeed, reads mapping to the beginning of the gRNA and sgRNA often presented an additional guanosine at the 5’ end, likely reflecting binding to the cap structure. This agrees with previous data reporting that GEMIN5 is a cap-binding protein (Bradrick and Gromeier, 2009; Xu et al., 2016). Additional peaks overlap with the downstream loop (DLP), which is a hairpin structure that stimulates the translation of the sgRNA (Frolov and Schlesinger, 1996). Interaction with the cap, 5’ UTR, 3’ UTR and DLP of viral RNAs aligns well with the proposed role as translational regulator and the observed inhibition of C expression. Strikingly, another GEMIN5 binding peak overlapped with the package signal recognised by SINV C, suggesting a potential interference with morphogenesis of viral particles. Our data strongly supports the model in which GEMIN5 recognises different regulatory regions of the viral RNA and prevents translation by interfering with ribosomal function.

Taken together, these results highlight the capacity of RNA-IC at identifying RBPs participating in SINV infection either as pro- or anti-viral factors.

## OUTLOOK

RNA-IC, and methods built on it, have been used in the past to compile the repertoire of cellular RBPs and their RBDs (Hentze et al., 2018). Nevertheless, we still poorly understand whether the RBPome globally responds to physiological cues and if so, how. Here, we show that the RBPome is highly dynamic and has the potential to adapt to the cellular state, in this case, infection with an RNA virus. SINV infection induced responses in a quarter of the identified RBPome, including both well-established and unconventional RBPs newly discovered by RNA-IC, evoking still unknown biological roles in infected cells. RBP dynamics highlight the post-transcriptional circuits modulated by the infection, which include the activation of the protein synthesis apparatus, the 5’ to 3’ RNA degradation machinery and a defined scope of antiviral factors as well as the inhibition of several RBPs involved in RNA processing and export. The differential RNA-binding activity of near ninety ribosomal and ribosome-associated proteins (Table S2) (Simsek et al., 2017), aligns well with the proposed, although not yet demonstrated, existence of specialised ribosomes in infected cells (Au and Jan, 2014). RNA helicases also emerge as responsive nodes in RBP networks.

Their RNP remodelling activity are linked to central regulatory roles in RNA metabolism and their ability to respond to biological cues likely contributes to their regulatory function. Both the existence of a specialised translation machinery and the role of dynamic RNA helicases in the post-transcriptional regulation of gene expression deserve further investigation.

Mechanistically, RBPome response to SINV infection is mostly driven by both deep alterations of the transcriptome and a global subcellular redistribution of relevant RBPs. In particular, a large proportion of stimulated RBPs are relocalised to the viral replication factories. Apart from these major alterations, RBP activity can be modulated in individual basis by more subtle mechanisms. For example, it is known that virus infection triggers signalling pathways involving kinases and E3 ubiquitin ligases (Figure 1D) (Gack et al., 2007; Greenwood et al., 2016; Li et al., 2017; Ventoso et al., 2006). This idea is strengthen here by the increased RNA-binding activity of the kinase SRPK1 and the E3 ubiquitin ligases TRIM25 and TRIM56. Hence, it is plausible that post-translational modifications also contribute to RBP regulation in SINV-infected cells. Furthermore, the interaction of GEMIN5 with viral proteins suggests that differential complex assembly might also regulate RBP activity (Figure 7C).

Importantly, RBP responses are biologically important as perturbation of dynamic RBPs strongly affect either the viral fitness or the capacity of the cell to counteract the infection. Therefore, every protein reported here to respond to SINV infection may offer new avenues of research due to their potential as anti- or pro-viral factors. We believe that these regulatory RBPs will represent excellent targets for novel therapeutic approaches. Moreover, viruses hijack master regulators of cellular pathways (Berget et al., 1977; Castello et al., 2011; Castello et al., 2012; Chow et al., 1977; Krausslich et al., 1987; Lloyd, 2015; Sun and Baltimore, 1989; von Kobbe et al., 2000), suggesting that responsive RBPs are likely involved in regulatory steps of RNA metabolism.

Some of the outstanding questions derived from this work include why the lack of the exonuclease XRN1 makes the cells refractory to SINV, what triggers the degradation of host RNA and why the transcripts induced by the antiviral response are resistant to degradation. Moreover, GEMIN5 emerges as a highly responsive RBP that impairs SINV infection. The extra-SMN complex activity of GEMIN5 in translational control has recently been reported (Francisco-Velilla et al., 2016; Pineiro et al., 2015). We show here that this newly discovered activity can be regulated, as GEMIN5 display altered RNA-binding activity and localisation in infected cells as it engages in interactions with viral proteins and RNAs. The exact mechanisms underpinning GEMIN5 effects in translation requires further investigation. Finally, ‘comparative RNA-IC’ has been, here, applied to cells infected with SINV. However, it can now be extended to other viruses or physiological cues to improve our understanding on RBP dynamics and its biological importance.

## AUTHOR CONTRIBUTIONS

Conceptualization, M.N., L.C., B.F., S.M., A.C.; Methodology, M.G.M, M.N., N.S., A.I.J, E.G.A, C.L., B.F., S.M., A.C.; Investigation, M.G.M., M.N., N.S., A.I.J., E.G.A, C.L., M.B.P, V.C., R.A., T.D., S.H., T.J.M.S., B.L., M.A.S., E.P.R., V.P., B.F., S.M., A.C.; Writing, original draft, M.G.M., M.N., A.I.J, A.C.; Writing, editing, M.G.M., M.N., N.S. A.I.J, C.L., S.M., A.C.; Funding acquisition, M.G.M, A.C.; Resources, L.C., E.P.R., V.P., B.F., S.M., A.C.; Supervision, M.G.M, M.N., A.I.J., B.F., S.M., A.C.

## ACKNOWLEDGEMENTS

We dedicate this work to our colleagues and friends Bernd Fischer and Katrin Eichelbaum, who sadly passed away during the development of this work. We thank Matthias W Hentze, Encarnacion Martinez-Salas, Gracjan Michlewski, Javier Martinez, Kui Li, Ilan Davis, Richard Parton and Jan Rehwinkel for reagents and advice. The support from Oxford Micron Advanced Bioimaging Unit in microscopy experiments is also acknowledged. A.C. is funded by MRC Career Development Award #MR/L019434/1 and the John Fell Funds from the University of Oxford. M.G.M is funded by the European Union’s Horizon 2020 research and innovation programme under the Marie-Sklodowska-Curie grant agreement No. 700184. V.P is funded by a SciLifeLab Fellowship, the Swedish Research Council (VR 2016-01842), a Wallenberg Academy Fellowship (KAW 2016.0123) and the Ragnar Söderberg Foundation. L.C. is funded by grant DGICYT SAF2015-66170-R (MINECO/FEDER). E.G.A. was awarded with a short-term EMBO fellowship (ASTF 358-2015) and the FPU Fellowship FPU15/05709.

## REFERENCES

Akhrymuk, I., Kulemzin, S.V., and Frolova, E.I. (2012). Evasion of the innate immune response: the Old World alphavirus nsP2 protein induces rapid degradation of Rpb1, a catalytic subunit of RNA polymerase II. J Virol 86, 7180–7191.

Au, H.H., and Jan, E. (2014). Novel viral translation strategies. Wiley Interdiscip Rev RNA 5, 779–801.

Balistreri, G., Horvath, P., Schweingruber, C., Zund, D., McInerney, G., Merits, A., Muhlemann, O., Azzalin, C., and Helenius, A. (2014). The host nonsense-mediated mRNA decay pathway restricts Mammalian RNA virus replication. Cell Host Microbe 16, 403–411.

Baltz, A.G., Munschauer, M., Schwanhausser, B., Vasile, A., Murakawa, Y., Schueler, M., Youngs, N., Penfold-Brown, D., Drew, K., Milek, M., et al. (2012). The mRNA-Bound Proteome and Its Global Occupancy Profile on Protein-Coding Transcripts. Mol Cell 46, 674–690.

Barbalat, R., Ewald, S.E., Mouchess, M.L., and Barton, G.M. (2011). Nucleic acid recognition by the innate immune system. Annu Rev Immunol 29, 185–214.

Battle, D.J., Kasim, M., Wang, J., and Dreyfuss, G. (2007). SMN-independent subunits of the SMN complex. Identification of a small nuclear ribonucleoprotein assembly intermediate. J Biol Chem 282, 27953–27959.

Berget, S.M., Moore, C., and Sharp, P.A. (1977). Spliced segments at the 5′ terminus of adenovirus 2 late mRNA. Proc Natl Acad Sci U S A 74, 3171–3175.

Bradrick, S.S., and Gromeier, M. (2009). Identification of gemin5 as a novel 7-methylguanosine cap-binding protein. PLoS One 4, e7030.

Brannan, K.W., Jin, W., Huelga, S.C., Banks, C.A., Gilmore, J.M., Florens, L., Washburn, M.P., Van Nostrand, E.L., Pratt, G.A., Schwinn, M.K., et al. (2016). SONAR Discovers RNA-Binding Proteins from Analysis of Large-Scale Protein-Protein Interactomes. Mol Cell 64, 282–293.

Carrasco, L., Sanz, M.A., and Gonzalez-Almela, E. (2018). The Regulation of Translation in Alphavirus-Infected Cells. Viruses 10.

Castello, A., Alvarez, E., and Carrasco, L. (2011). The multifaceted poliovirus 2A protease: regulation of gene expression by picornavirus proteases. J Biomed Biotechnol 2011, 369648.

Castello, A., Fischer, B., Eichelbaum, K., Horos, R., Beckmann, B.M., Strein, C., Davey, N.E., Humphreys, D.T., Preiss, T., Steinmetz, L.M., et al. (2012). Insights into RNA Biology from an Atlas of Mammalian mRNA-Binding Proteins. Cell 149, 1393–1406.

Castello, A., Fischer, B., Frese, C.K., Horos, R., Alleaume, A.M., Foehr, S., Curk, T., Krijgsveld, J., and Hentze, M.W. (2016). Comprehensive Identification of RNA-Binding Domains in Human Cells. Mol Cell 63, 696–710.

Castello, A., Horos, R., Strein, C., Fischer, B., Eichelbaum, K., Steinmetz, L.M., Krijgsveld, J., and Hentze, M.W. (2013). System-wide identification of RNA-binding proteins by interactome capture. Nat Protoc 8, 491–500.

Castello, A., Sanz, M.A., Molina, S., and Carrasco, L. (2006). Translation of Sindbis virus 26S mRNA does not require intact eukariotic initiation factor 4G. J Mol Biol 355, 942–956.

Chang, C.H., Curtis, J.D., Maggi, L.B., Jr., Faubert, B., Villarino, A.V., O’Sullivan, D., Huang, S.C., van der Windt, G.J., Blagih, J., Qiu, J., et al. (2013). Posttranscriptional control of T cell effector function by aerobic glycolysis. Cell 153, 1239–1251.

Chen, C.Y., and Shyu, A.B. (2014). Emerging mechanisms of mRNP remodeling regulation. Wiley Interdiscip Rev RNA 5, 713–722.

Choudhury, N.R., Heikel, G., Trubitsyna, M., Kubik, P., Nowak, J.S., Webb, S., Granneman, S., Spanos, C., Rappsilber, J., Castello, A., et al. (2017). RNA-binding activity of TRIM25 is mediated by its PRY/SPRY domain and is required for ubiquitination. BMC Biol 15, 105.

Chow, L.T., Gelinas, R.E., Broker, T.R., and Roberts, R.J. (1977). An amazing sequence arrangement at the 5′ ends of adenovirus 2 messenger RNA. Cell 12, 1–8.

Francisco-Velilla, R., Fernandez-Chamorro, J., Ramajo, J., and Martinez-Salas, E. (2016). The RNA-binding protein Gemin5 binds directly to the ribosome and regulates global translation. Nucleic Acids Res 44, 8335–8351.

Frolov, I., and Schlesinger, S. (1996). Translation of Sindbis virus mRNA: analysis of sequences downstream of the initiating AUG codon that enhance translation. J Virol 70, 1182–1190.

Fukuhara, T., Hosoya, T., Shimizu, S., Sumi, K., Oshiro, T., Yoshinaka, Y., Suzuki, M., Yamamoto, N., Herzenberg, L.A., Herzenberg, L.A., et al. (2006). Utilization of host SR protein kinases and RNA-splicing machinery during viral replication. Proc Natl Acad Sci U S A 103, 11329–11333.

Gack, M.U., Shin, Y.C., Joo, C.H., Urano, T., Liang, C., Sun, L., Takeuchi, O., Akira, S., Chen, Z., Inoue, S., et al. (2007). TRIM25 RING-finger E3 ubiquitin ligase is essential for RIG-I-mediated antiviral activity. Nature 446, 916–920.

Garcia-Moreno, M., Jaervelin, A.I., and Castello, A. (2018). Unconventional RNA-binding proteins step into the virus-host battlefront. WIREs RNA In press.

Garcia-Moreno, M., Sanz, M.A., and Carrasco, L. (2015). Initiation codon selection is accomplished by a scanning mechanism without crucial initiation factors in Sindbis virus subgenomic mRNA. RNA 21, 93–112.

Garcia-Moreno, M., Sanz, M.A., Pelletier, J., and Carrasco, L. (2013). Requirements for eIF4A and eIF2 during translation of Sindbis virus subgenomic mRNA in vertebrate and invertebrate host cells. Cell Microbiol 15, 823–840.

Garcia, M.A., Gil, J., Ventoso, I., Guerra, S., Domingo, E., Rivas, C., and Esteban, M. (2006). Impact of protein kinase PKR in cell biology: from antiviral to antiproliferative action. Microbiol Mol Biol Rev 70, 1032–1060.

Geller, R., Taguwa, S., and Frydman, J. (2012). Broad action of Hsp90 as a host chaperone required for viral replication. Bba-Mol Cell Res 1823, 698–706.

Gerstberger, S., Hafner, M., and Tuschl, T. (2014). A census of human RNA-binding proteins. Nat Rev Genet 15, 829–845.

Glisovic, T., Bachorik, J.L., Yong, J., and Dreyfuss, G. (2008). RNA-binding proteins and post-transcriptional gene regulation. FEBS Lett 582, 1977–1986.

Goodier, J.L., Cheung, L.E., and Kazazian, H.H., Jr. (2012). MOV10 RNA helicase is a potent inhibitor of retrotransposition in cells. PLoS Genet 8, e1002941.

Gorchakov, R., Frolova, E., and Frolov, I. (2005). Inhibition of transcription and translation in Sindbis virus-infected cells. J Virol 79, 9397–9409.

Greenwood, E.J., Matheson, N.J., Wals, K., van den Boomen, D.J., Antrobus, R., Williamson, J.C., and Lehner, P.J. (2016). Temporal proteomic analysis of HIV infection reveals remodelling of the host phosphoproteome by lentiviral Vif variants. eLife 5.

Gubitz, A.K., Mourelatos, Z., Abel, L., Rappsilber, J., Mann, M., and Dreyfuss, G. (2002). Gemin5, a novel WD repeat protein component of the SMN complex that binds Sm proteins. J Biol Chem 277, 5631–5636.

He, C., Sidoli, S., Warneford-Thomson, R., Tatomer, D.C., Wilusz, J.E., Garcia, B.A., and Bonasio, R. (2016). High-Resolution Mapping of RNA-Binding Regions in the Nuclear Proteome of Embryonic Stem Cells. Mol Cell 64, 416–430.

Hentze, M.W., and Argos, P. (1991). Homology between IRE-BP, a regulatory RNA-binding protein, aconitase, and isopropylmalate isomerase. Nucleic Acids Res 19, 1739–1740.

Hentze, M.W., Castello, A., Schwarzl, T., and Preiss, T. (2018). A brave new world of RNA-binding proteins. Nat Rev Mol Cell Biol.

Howard, J.M., and Sanford, J.R. (2015). The RNAissance family: SR proteins as multifaceted regulators of gene expression. Wiley Interdiscip Rev RNA 6, 93–110.

Iwasaki, S., Kobayashi, M., Yoda, M., Sakaguchi, Y., Katsuma, S., Suzuki, T., and Tomari, Y. (2010). Hsc70/Hsp90 chaperone machinery mediates ATP-dependent RISC loading of small RNA duplexes. Mol Cell 39, 292–299.

Jurkin, J., Henkel, T., Nielsen, A.F., Minnich, M., Popow, J., Kaufmann, T., Heindl, K., Hoffmann, T., Busslinger, M., and Martinez, J. (2014). The mammalian tRNA ligase complex mediates splicing of XBP1 mRNA and controls antibody secretion in plasma cells. EMBO J 33, 2922–2936.

König, J., Zarnack, K., Rot, G., Curk, T., Kayikci, M., Zupan, B., Turner, D.J., Luscombe, N.M., and Ule, J. (2010). iCLIP reveals the function of hnRNP particles in splicing at individual nucleotide resolution. Nat Struct Mol Biol 17, 909–915.

Kramer, K., Sachsenberg, T., Beckmann, B.M., Qamar, S., Boon, K.L., Hentze, M.W., Kohlbacher, O., and Urlaub, H. (2014). Photo-cross-linking and high-resolution mass spectrometry for assignment of RNA-binding sites in RNA-binding proteins. Nat Methods 11, 1064–1070.

Krausslich, H.G., Nicklin, M.J., Toyoda, H., Etchison, D., and Wimmer, E. (1987). Poliovirus proteinase 2A induces cleavage of eucaryotic initiation factor 4F polypeptide p220. J Virol 61, 2711–2718.

LaPointe, A.T., Gebhart, N.N., Meller, M.E., Hardy, R.W., and Sokoloski, K.J. (2018). The Identification and Characterization of Sindbis Virus RNA:Host Protein Interactions. J Virol.

Lee, A.S., Kranzusch, P.J., Doudna, J.A., and Cate, J.H. (2016). eIF3d is an mRNA cap-binding protein that is required for specialized translation initiation. Nature 536, 96–99.

Li, J., Tang, S., Hewlett, I., and Yang, M. (2007). HIV-1 capsid protein and cyclophilin a as new targets for anti-AIDS therapeutic agents. Infect Disord Drug Targets 7, 238–244.

Li, M.M., Lau, Z., Cheung, P., Aguilar, E.G., Schneider, W.M., Bozzacco, L., Molina, H., Buehler, E., Takaoka, A., Rice, C.M., et al. (2017). TRIM25 Enhances the Antiviral Action of Zinc-Finger Antiviral Protein (ZAP). PLoS Pathog 13, e1006145.

Lin, J.Y., Shih, S.R., Pan, M., Li, C., Lue, C.F., Stollar, V., and Li, M.L. (2009). hnRNP A1 interacts with the 5′ untranslated regions of enterovirus 71 and Sindbis virus RNA and is required for viral replication. J Virol 83, 6106–6114.

Liu, C., Perilla, J.R., Ning, J., Lu, M., Hou, G., Ramalho, R., Himes, B.A., Zhao, G., Bedwell, G.J., Byeon, I.J., et al. (2016). Cyclophilin A stabilizes the HIV-1 capsid through a novel non-canonical binding site. Nat Commun 7, 10714.

Lloyd, R.E. (2015). Nuclear proteins hijacked by mammalian cytoplasmic plus strand RNA viruses. Virology 479-480, 457–474.

Lunde, B.M., Moore, C., and Varani, G. (2007). RNA-binding proteins: modular design for efficient function. Nat Rev Mol Cell Biol 8, 479–490.

Manokaran, G., Finol, E., Wang, C., Gunaratne, J., Bahl, J., Ong, E.Z., Tan, H.C., Sessions, O.M., Ward, A.M., Gubler, D.J., et al. (2015). Dengue subgenomic RNA binds TRIM25 to inhibit interferon expression for epidemiological fitness. Science 350, 217–221.

Mesa, A., Somarelli, J.A., and Herrera, R.J. (2008). Spliceosomal immunophilins. FEBS Lett 582, 2345–2351.

Molleston, J.M., and Cherry, S. (2017). Attacked from All Sides: RNA Decay in Antiviral Defense. Viruses 9.

Monie, T.P., Perrin, A.J., Birtley, J.R., Sweeney, T.R., Karakasiliotis, I., Chaudhry, Y., Roberts, L.O., Matthews, S., Goodfellow, I.G., and Curry, S. (2007). Structural insights into the transcriptional and translational roles of Ebp1. EMBO J 26, 3936–3944.

Mukherjee, N., Calviello, L., Hirsekorn, A., de Pretis, S., Pelizzola, M., and Ohler, U. (2017). Integrative classification of human coding and noncoding genes through RNA metabolism profiles. Nat Struct Mol Biol 24, 86–96.

Ni, X., Ru, H., Ma, F., Zhao, L., Shaw, N., Feng, Y., Ding, W., Gong, W., Wang, Q., Ouyang, S., et al. (2016). New insights into the structural basis of DNA recognition by HINa and HINb domains of IFI16. J Mol Cell Biol 8, 51–61.

Ong, S.E., and Mann, M. (2006). A practical recipe for stable isotope labeling by amino acids in cell culture (SILAC). Nat Protoc 1, 2650–2660.

Pacheco, A., Lopez de Quinto, S., Ramajo, J., Fernandez, N., and Martinez-Salas, E. (2009). A novel role for Gemin5 in mRNA translation. Nucleic Acids Res 37, 582–590.

Pineiro, D., Fernandez-Chamorro, J., Francisco-Velilla, R., and Martinez-Salas, E. (2015). Gemin5: A Multitasking RNA-Binding Protein Involved in Translation Control. Biomolecules 5, 528–544.

Popow, J., Englert, M., Weitzer, S., Schleiffer, A., Mierzwa, B., Mechtler, K., Trowitzsch, S., Will, C.L., Luhrmann, R., Soll, D., et al. (2011). HSPC117 is the essential subunit of a human tRNA splicing ligase complex. Science 331, 760–764.

Popow, J., Jurkin, J., Schleiffer, A., and Martinez, J. (2014). Analysis of orthologous groups reveals archease and DDX1 as tRNA splicing factors. Nature 511, 104–107.

Rathore, A.P., Ng, M.L., and Vasudevan, S.G. (2013). Differential unfolded protein response during Chikungunya and Sindbis virus infection: CHIKV nsP4 suppresses eIF2alpha phosphorylation. Virol J 10, 36.

Rehwinkel, J., and Reis e Sousa, C. (2013). Targeting the viral Achilles’ heel: recognition of 5′-triphosphate RNA in innate anti-viral defence. Curr Opin Microbiol 16, 485–492.

Romero-Brey, I., and Bartenschlager, R. (2014). Membranous replication factories induced by plus-strand RNA viruses. Viruses 6, 2826–2857.

Rupp, D., and Bartenschlager, R. (2014). Targets for antiviral therapy of hepatitis C. Semin Liver Dis 34, 9–21.

Sampath, P., Mazumder, B., Seshadri, V., Gerber, C.A., Chavatte, L., Kinter, M., Ting, S.M., Dignam, J.D., Kim, S., Driscoll, D.M., et al. (2004). Noncanonical function of glutamyl-prolyl-tRNA synthetase: gene-specific silencing of translation. Cell 119, 195–208.

Seo, G.J., Kim, C., Shin, W.J., Sklan, E.H., Eoh, H., and Jung, J.U. (2018). TRIM56-mediated monoubiquitination of cGAS for cytosolic DNA sensing. Nat Commun 9, 613.

Shen, Y., Li, N.L., Wang, J., Liu, B., Lester, S., and Li, K. (2012). TRIM56 is an essential component of the TLR3 antiviral signaling pathway. J Biol Chem 287, 36404–36413.

Simsek, D., Tiu, G.C., Flynn, R.A., Byeon, G.W., Leppek, K., Xu, A.F., Chang, H.Y., and Barna, M. (2017). The Mammalian Ribo-interactome Reveals Ribosome Functional Diversity and Heterogeneity. Cell 169, 1051–1065 e1018.

Skabkin, M.A., Skabkina, O.V., Dhote, V., Komar, A.A., Hellen, C.U., and Pestova, T.V. (2010). Activities of Ligatin and MCT-1/DENR in eukaryotic translation initiation and ribosomal recycling. Genes Dev 24, 1787–1801.

Sun, X.H., and Baltimore, D. (1989). Human immunodeficiency virus tat-activated expression of poliovirus protein 2A inhibits mRNA translation. Proc Natl Acad Sci U S A 86, 2143–2146.

Sysoev, V.O., Fischer, B., Frese, C.K., Gupta, I., Krijgsveld, J., Hentze, M.W., Castello, A., and Ephrussi, A. (2016). Global changes of the RNA-bound proteome during the maternal-to-zygotic transition in Drosophila. Nat Commun 7, 12128.

Thompson, M.R., Sharma, S., Atianand, M., Jensen, S.B., Carpenter, S., Knipe, D.M., Fitzgerald, K.A., and Kurt-Jones, E.A. (2014). Interferon gamma-inducible protein (IFI) 16 transcriptionally regulates type i interferons and other interferon-stimulated genes and controls the interferon response to both DNA and RNA viruses. J Biol Chem 289, 23568–23581.

Tsuchida, T., Zou, J., Saitoh, T., Kumar, H., Abe, T., Matsuura, Y., Kawai, T., and Akira, S. (2010). The ubiquitin ligase TRIM56 regulates innate immune responses to intracellular double-stranded DNA. Immunity 33, 765–776.

Ventoso, I., Sanz, M.A., Molina, S., Berlanga, J.J., Carrasco, L., and Esteban, M. (2006). Translational resistance of late alphavirus mRNA to eIF2alpha phosphorylation: a strategy to overcome the antiviral effect of protein kinase PKR. Genes Dev 20, 87–100.

Versteeg, G.A., Rajsbaum, R., Sanchez-Aparicio, M.T., Maestre, A.M., Valdiviezo, J., Shi, M., Inn, K.S., Fernandez-Sesma, A., Jung, J., and Garcia-Sastre, A. (2013). The E3-ligase TRIM family of proteins regulates signaling pathways triggered by innate immune pattern-recognition receptors. Immunity 38, 384–398.

Vladimer, G.I., Gorna, M.W., and Superti-Furga, G. (2014). IFITs: Emerging Roles as Key Anti-Viral Proteins. Front Immunol 5, 94.

von Kobbe, C., van Deursen, J.M., Rodrigues, J.P., Sitterlin, D., Bachi, A., Wu, X., Wilm, M., Carmo-Fonseca, M., and Izaurralde, E. (2000). Vesicular stomatitis virus matrix protein inhibits host cell gene expression by targeting the nucleoporin Nup98. Mol Cell 6, 1243–1252.

Wang, Y., Jin, F., Wang, R., Li, F., Wu, Y., Kitazato, K., and Wang, Y. (2017). HSP90: a promising broad-spectrum antiviral drug target. Arch Virol 162, 3269–3282.

Xu, C., Ishikawa, H., Izumikawa, K., Li, L., He, H., Nobe, Y., Yamauchi, Y., Shahjee, H.M., Wu, X.H., Yu, Y.T., et al. (2016). Structural insights into Gemin5-guided selection of pre-snRNAs for snRNP assembly. Genes Dev 30, 2376–2390.

Zhang, R., Hryc, C.F., Cong, Y., Liu, X., Jakana, J., Gorchakov, R., Baker, M.L., Weaver, S.C., and Chiu, W. (2011). 4.4 A cryo-EM structure of an enveloped alphavirus Venezuelan equine encephalitis virus. EMBO J 30, 3854–3863.

Zheng, X., Wang, X., Tu, F., Wang, Q., Fan, Z., and Gao, G. (2017). TRIM25 Is Required for the Antiviral Activity of Zinc Finger Antiviral Protein. J Virol 91.

Zhou, D., Mei, Q., Li, J., and He, H. (2012). Cyclophilin A and viral infections. Biochem Biophys Res Commun 424, 647–650.

